# Involvement of the medial prefrontal cortex in food addiction: new insights from *in vivo* calcium Imaging

**DOI:** 10.64898/2025.12.10.693384

**Authors:** Roberto Capellán, Ignacio Marin-Blasco, Carles Ibañez, Lucas Perez-Molina, Johans Pacheco, Raul Andero, Rafael Maldonado, Elena Martín-García

## Abstract

Food addiction is a multifactorial disorder characterized by a loss of control over food intake, contributing to the development of obesity. To investigate the involvement of the prelimbic (PL) medial prefrontal cortex (mPFC) in the transition to food addiction, we examined PL calcium dynamics in mice exhibiting addicted versus non-addicted behavioral phenotypes. Our behavioral results allowed the identification of two distinct subpopulations of addicted and non-addicted mice, enabling direct comparison of their neural activity profiles. Addicted and non-addicted mice showed significant differences across several PL calcium activity parameters. These findings demonstrate a strong association between PL mPFC calcium activity dynamics and key addiction criteria, highlighting a critical role of this brain region in the development of food addiction. Understanding these neurobiological differences enhances our insight into brain mechanisms underlying loss of eating control and may inform more targeted approaches for studying and ultimately treating food addiction.

## INTRODUCTION

Food addiction is a controversial and still-debated concept [1], described as a complex, multifactorial behavioral disorder marked by a loss of control over food intake and a rising prevalence in recent years [2]. It is characterized by compulsive consumption and loss of control of palatable food intake, which induces adaptive changes in the brain’s reward circuitry. This maladaptive behavior is associated with obesity and other eating disorders and currently lacks effective treatments, contributing to substantial socioeconomic costs worldwide [3]. Although the concept of food addiction was proposed decades ago [4], it is not included in the 5th edition of the Diagnostic and Statistical Manual of Mental Disorders (DSM-5) [5]. Nevertheless, a widely used clinical tool to assess food addiction is the Yale Food Addiction Scale (YFAS), updated in 2016 to YFAS 2.0 to align with DSM-5 substance use disorder criteria [6]. The YFAS 2.0 criteria can be summarized into three hallmarks—persistent food seeking, heightened motivation to obtain food, and compulsive-like behavior—, which are also commonly used to model this disorder in rodents [7,8].

In previous work, we validated a mouse model of food addiction [9], which has been also later used and replicated by other reserchers [7,8]. We identified a specific prelimbic (PL) medial prefrontal cortex (mPFC)–nucleus accumbens (NA) circuit underlying vulnerability to developing food addiction and described associated epigenetic mechanisms contributing to this multifactorial pathology [7,8]. More recently, we characterized extreme subpopulations of food-addicted and non-addicted mice and identified distinct gut microbiota signatures linked to food addiction vulnerability. Indeed, we applied the YFAS 2.0 using parallel behavioral criteria to a human cohort and found corresponding gut microbiota signatures that may serve as potential biomarkers [10]. We further demonstrated the functional relevance of *Blautia*—the genus most differentially expressed in both addicted mice and humans—by administering the non-digestible carbohydrates lactulose and rhamnose [11,12], which increased gut *Blautia* abundance and prevented the development of food addiction in mice.

In the present study, we investigated the role of the PL mPFC in the transition to food addiction by examining PL calcium dynamics in mice displaying addicted versus non-addicted behavioral phenotypes. Understanding these neurobiological distinctions deepens our insight into the mechanisms underlying loss of eating control and may inform more targeted strategies for studying and ultimately treating food addiction.

## METHODS

### 1. Animals

Fifteen male C57BL/6J mice, aged eight weeks (Charles River, Lyon, France), were housed individually under controlled laboratory conditions (21 ± 1 °C; 55 ± 10% humidity) with *ad libitum* access to food and water throughout the experiment. Behavioural testing was conducted during the dark phase of a reversed light cycle (lights off at 7:30 a.m. and on at 7:30 p.m.). All procedures were performed in strict accordance with the European Communities Council Directive 2010/63/EU and were approved by the local ethical committee (Comitè Ètic d’Experimentació Animal–Parc de Recerca Biomèdica de Barcelona, CEEAPRBB, Protocol Number: RML-20-0053-P1) and the Generalitat de Catalunya (Protocol Number: DAAM-9687). The animal facilities operate under Animal Welfare Assurance (#A5388-01, Institutional Animal Care and Use Committee approval date: 05/08/2009) granted by the Office of Laboratory Animal Welfare (OLAW), U.S. National Institutes of Health.

### 2. Stereotaxic surgery and histological validation

#### 2.1 Viral injections

Animals were anesthetized with isoflurane (IsoFlo, Proima Ganadera SL, Spain) in oxygen at 4% for induction, and 2.0–2.5% for maintenance, with an airflow of 1.25 L/ min. The head of the animal were correctly placed in the stereotaxic frame (Kopf Model 962, Harvard-Panlab, Barcelona, Spain), according to bregma and lambda. The skull was exposed and cleaned using saline serum (0.9% NaCl). Animals were then injected with 800 nL of AAV expressing GCaMP6f calcium indicator (AAV1/2-Syn-WPRE-SV40-GCaMP6f; Addgene, USA) in the following coordinates (AP + 1.8 mm, LM ± 0.3 mm, DV – 2.3 mm) in a rate of 0.07 μL/min using a Hamilton syringe (75RN model; Cibertec-Harvard, Madrid, Spain) coupled to an injection pump (KD Scientific, MA, USA). After the injection, the syringe was slowly removed from the brain, ensuring that no liquid refluxed from the injection site. Animals were then sutured and left to rest under close supervision for one week.

#### 2.1 GRIN lens implantation and baseplating

After one week recovery period, animals were anesthetized and head-fixed in a stereotaxic frame. After exposing the skull, a 0.5 mm-tip drill bit was used to perforate the bone at the previously mentioned AP and LM coordinates. Then, a guide hole was made into the brain using the stereotaxic device and a 0.5 × 16 mm blunt syringe needle, reaching a depth of 100 μm above the viral injection site (DV -2.2 mm). Then, a 1.0 pitch, 0.5 mm-diameter, 4 mm-length GRIN lens (Inscopix, CA, USA), was implanted precisely at the same AAV injection site (DV -2.3 mm). Throughout this process, saline serum was constantly applied to the entry point as a lubricant. After complete insertion, the lenses were fixed in place with cyanoacrylate adhesive. Two screws were implanted on the skull surface as anchoring system. Then, the exposed scalp, lens, and screws were covered and secured with dental cement (Rebaron, GC Dental), and the area was stained with a small amount of black ink (Edding T-100). The exposed sections of the lenses were then protected with plastic caps (0.2 mL PCR Eppendorf tube caps) fixed with cyanoacrylate adhesive to the dental cement. Animals were individually housed to prevent complications associated with the implants due to social contact and were monitored during a 15-day recovery. After two weeks of post-operative care, the protective caps were removed, and a metallic baseplate was attached to the skull using stained dental cement. The calcium signal was simultaneously monitored with the Miniscope to ensure the optimal optical focus for each animal. Animals were then left to rest for one week before starting the behavioral test.

### 3. Food addiction-like behaviour assessment

Mice were trained to obtain chocolate-flavoured pellets in operant conditioning chambers during 24 daily 1-h self-administration sessions. Operant responding maintained by food was conducted in mouse operant chambers (Model ENV-307A-CT, Med Associates, Georgia, VT, USA) equipped with two retractable levers, one designated as the active lever and the other as the inactive lever. The chambers, constructed of aluminium and acrylic, were enclosed in sound- and light-attenuating cubicles containing ventilation fans that provided white noise. A food dispenser, positioned equidistant between the two levers, delivered pellets as required. The grid floor consisted of parallel metal bars capable of delivering electric foot shocks when scheduled. Presses on the active lever resulted in pellet delivery accompanied by an illuminated cue light positioned above the lever, whereas responses on the inactive lever had no programmed consequences.

Mice first completed six fixed-ratio 1 (FR1) sessions, followed by eighteen fixed-ratio 5 (FR5) sessions (Figure 1). Throughout this short operant conditioning program maintained by chocolate-flavoured pellets, three food addiction–like criteria were assessed: persistence of response, motivation, and compulsive-like behaviour [8,9,13]. Persistence of response was evaluated across three consecutive days following the progressive ratio (PR) session by quantifying non-reinforced active responses during a 10-min pellet-free period in which the chamber was illuminated, signalling pellet unavailability. Motivation was assessed using a PR schedule in which the response requirement to earn a pellet increased according to the following series: 1, 5, 12, 21, 33, 51, 75, 90, 120, 155, 180, 225, 260, 300, 350, 410, 465, 540, 630, 730, 850, 1000, 1200, 1500, 1800, 2100, 2400, 2700, 3000, 3400, 3800, 4200, 4600, 5000, and 5500. The breaking point was defined as the last ratio at which a pellet was obtained. Sessions lasted up to 5 h or ended if animals ceased responding on both levers for 1 h. Compulsive-like behaviour was determined in a 50-min FR5 session in which each pellet delivery was paired with punishment. Each earned pellet was associated with a 0.13 mA, 2-s foot shock. On the 4th active-lever response, mice received a shock without pellet delivery; on the 5th response, they received a second shock, accompanied by pellet delivery and the cue light. The schedule resets 10 sec after pellet delivery (time-out) or after the 4th response if the 5th response was not completed within 1 min.

**Figure 1.**
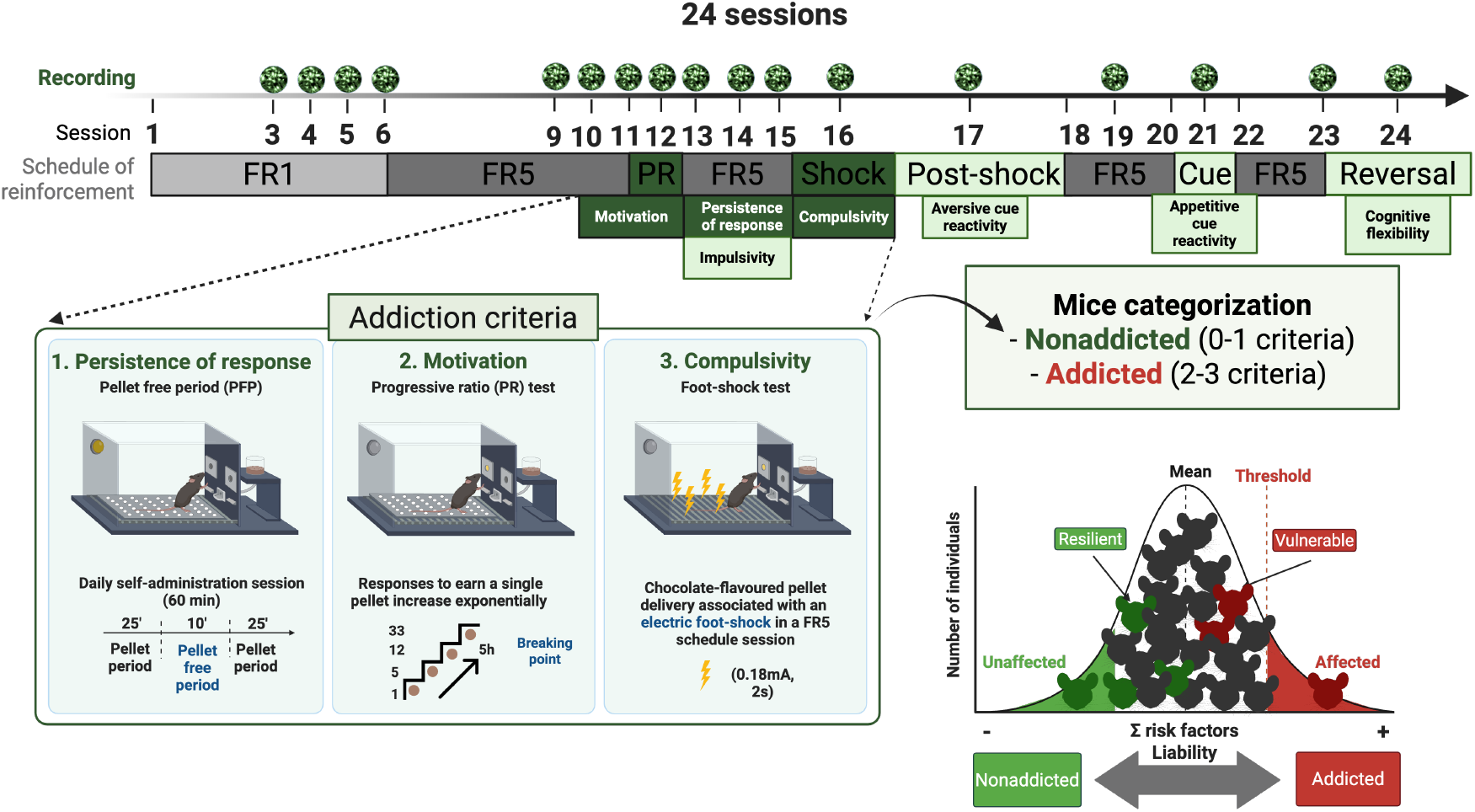
Assessment of food addiction-like behaviour. Experimental timeline of operant conditioning sessions; icons above specific sessions indicate days on which calcium imaging recordings were performed.

These behavioural criteria reflect key features of human food addiction as evaluated by the Yale Food Addiction Scale (YFAS 2.0), based on DSM-5 criteria for substance use disorders. Mice were considered positive for an addiction-like criterion when their performance met or exceeded the 75th percentile of the control-group distribution. Animals achieving two or three criteria were classified as addicted, whereas those meeting zero or one criterion were classified as non-addicted behaviour [7,9,13].

Additionally, four phenotypic traits associated with vulnerability to addiction were examined to obtain a comprehensive addiction-like behavioural profile: motor impulsivity, cognitive inflexibility, aversive associative learning, and appetitive associative learning [13]. Motor impulsivity, defined as the failure to inhibit a response once initiated, was quantified as non-reinforced active responses during the 10-sec time-out period after each pellet delivery across the three days preceding the PR test [14]. Cognitive inflexibility was assessed through a reversal learning session in which the identities of the active and inactive levers were switched relative to previous sessions. Aversive associative learning was evaluated the day after the shock-paired session, using the same context but without delivering shocks, pellets or presenting the cue light; non-reinforced active responses were recorded as an index of aversive learning. Appetitive associative learning was measured in a 90-min cue-induced food-seeking test consisting of a 60-min extinction period in which lever presses were not reinforced, followed by a 30-min period during which the pellet-associated cue light was presented contingently under an FR5 schedule.

### 4. Calcium imaging recording and analysis

#### 4.1 General considerations

Calcium activity (GCaMP6f 488nm fluorescence) was registered using a Miniscope (v4.0 UCLA University, CA, USA) coupled to the Miniscope DAQ hardware and software. The Miniscope system is an open-source microscopy platform for recording and analyzing neural activity in freely behaving animals, developed by Daniel Aharoni Lab (UCLA University, CA, USA). The Miniscope DAQ software was configured to record at 20 frames per second. The LED intensity and digital gain were adjusted for each animal and kept constant throughout the experiment. In a subset of recordings, minor technical artefacts intrinsic to head-mounted calcium imaging (such as transient motion-related fluctuations, occasional LED intensity instability, or brief loss of focus) were detected during quality control. These segments were automatically flagged and excluded before analysis, ensuring that only stable, high-quality fluorescence traces contributed to event-locked neural measures. No systematic differences in recording quality were observed between experimental groups, and therefore these artefacts are unlikely to have biased the reported results.

#### 4.2 Calcium signal deconvolution

We used the publicly available algorithm, Constrained Non-Negative Matrix Factorisation optimized for Endoscope signals (CNMF-E), to deconvolve the calcium signal (Zhou et al 2018). CNMF-E first identifies non-linear shifts in the visual field using an optimized template-matching algorithm, NoRMCorre [15].

All data were reconstructed using the shift vectors obtained during motion registration, thereby removing frame-to-frame displacement artifacts. Following motion correction, the field of view was partitioned into spatial patches for processing with the CNMF-E algorithm. For each patch, a peak-to-noise ratio (PNR) relative to baseline fluorescence was computed together with a local temporal correlation metric. These measures were used to identify high-quality candidate pixels and delineate the spatial footprints of individual neuronal somata. Using these spatial and temporal features, CNMF-E detects, separates, and characterizes neuronal components, enabling the discrimination of partially overlapping neurons and the demixing of their corresponding temporal signals. The fluorescence trace associated with each region of interest was then fit with a spike-decay model consistent with the calcium indicator’s kinetics and converted to relative fluorescence change (ΔF/F).

For all sessions, the following set-up was used: spatial downsampling = 3; motion correction = non-rigid; temporal downsampling = 3; dendrite identification = false; spatial algorithm = hals; include residuals = false; deconvolution method = foopsi, minimum spike size = 5 x noise; de-trend method = spline; background model = svd.

To ensure the reliability of the extracted neuronal activity, a multi-stage filtering procedure was applied. Components exhibiting abnormally high peak amplitudes, values inconsistent with established calcium transient dynamics, were excluded using confidence-interval thresholds set at the 95th or 99th percentile, based on inspection of the amplitude distribution. Components with spatial footprints extending beyond the optical field of view were removed by applying a spatial mask. In a small subset of cases, remaining components underwent manual inspection to exclude traces whose temporal profiles were not consistent with biologically plausible neuronal activity. This curation procedure ensured that only high-confidence neuronal signals were retained for downstream analyses.

#### 4.3 PL calcium activity during specific events

Calcium dynamics were quantified for each behavioural hallmark of addiction-like phenotypes by temporally aligning fluorescence traces to the onset of discrete operant events, including active lever presses, inactive lever presses, footshock delivery, reward presentation, and conditioned cue delivery. To prevent photobleaching and maintain stable signal quality across long behavioural sessions, calcium imaging was acquired in predefined 5-min recording epochs embedded within each task. These structured epochs were positioned at key behavioural phases (early, intermediate, late, and event-specific periods such as pellet-free intervals, late breaking points, shock delivery, or cue presentation), providing consistent sampling across animals while minimizing signal degradation. Event-locked neural activity was analysed using a ±1 sec peri-event window and an additional 10 sec post-event window, from which mean fluorescence values were extracted to generate activity profiles associated with motivation, persistence of responding, compulsivity, impulsivity, aversive associative learning, and cognitive inflexibility. To enable reliable comparisons across animals, event-evoked fluorescence changes were normalized within each recording epoch. Specifically, the mean fluorescence measured in the ±1 sec peri-event window was expressed relative to the average fluorescence level of the 5-min recording block in which the event occurred. This normalization controlled for inter-individual variability in baseline calcium signal and ensured that analyses reflected relative event-driven changes rather than differences in absolute fluorescence magnitude.

#### 4.4 Identifying stimulus-responding neurons

Stimulus–response coupling was quantified by binarizing each neural trace to indicate the temporal occurrence of calcium events, defined as local maxima in the fluorescence signal. Behavioural event presence was encoded within a peri-event window spanning 5 sec before event onset and 10 sec following event onset.

For each neuron, the Phi coefficient (*ϕ*) (Equation 1) was computed to quantify the association between behavioural response presence and the occurrence of calcium events. True positives were defined as frames in which a calcium event occurred during a behavioural response. In contrast, false negatives corresponded to frames lacking a calcium event despite the presence of a behavioural response. To avoid imposing a global *ϕ* threshold and to control for spurious correlations arising from the intrinsic peak structure of each trace, event matrices were randomized while preserving the total number of calcium events per neuron. For each neuron, *ϕ* values were recomputed across 1000 (*m*) randomized iterations, and the resulting distribution was stored (Equation 2). The observed *ϕ* value was then converted to a Z-score (Equation 3) by subtracting the mean *ϕ* of the randomized distribution and dividing by its standard deviation (*σR*). Neurons were classified as active if their *ϕ* Z-score exceeded the upper confidence interval (Z ≥ 1.96) of the randomized distribution, indicating a significant positive association between calcium events and behavioural response presence. Conversely, inactive neurons were defined as those with *ϕ* Z-scores below the lower confidence interval (Z ≤ –1.96), indicating a reliable absence of calcium events during the peri-event behavioural window.

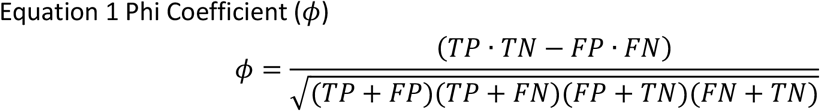

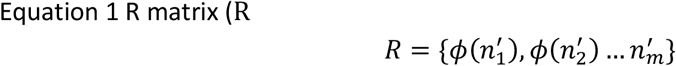

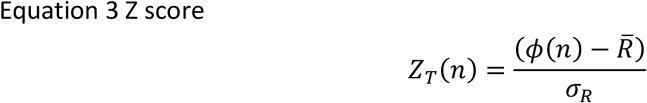

### 5. Histological processing and verification

After completion of the experimental protocol, animals were euthanized by rapid decapitation, and their brains were rapidly extracted. Brains were post-fixed in 4% paraformaldehyde for 24 h at 4 °C and subsequently transferred to 0.1 M phosphate buffer (PB) for storage until histological processing. PL coronal sections (30 μm) were obtained using a vibratome (Leica VT1000S). Sections were mounted on Superfrost slides (Thermo Scientific, USA), and fluorescence images were acquired using a Nikon Eclipse 90i microscope to visualize the distribution of the vector-associated reporter (EGFP). These images were used to verify the accuracy of the AAV injection sites and the placement of the imaging lens.

A mouse brain atlas image showing AAV vector spread and GRIN lens placement in the PL region was included in the supplementary material (Fig. S1).

### 6. Statistical analyses

All analyses were performed in IBM SPSS Statistics 27. Normality and homogeneity of variance were assessed using the Shapiro–Wilk and Levene tests, respectively. Parametric tests (independent-samples t-test, repeated-measures ANOVA, mixed-model ANOVA) were used when assumptions were met; otherwise, non-parametric tests (Mann–Whitney U) were applied. Post hoc comparisons were corrected using the Bonferroni method. Event-evoked calcium fluorescence was analysed using the same criteria, with group comparisons performed separately for each behavioural event type. All tests were two-tailed with α = 0.05, and results are presented as mean ± SEM.

## RESULTS

In the operant assessment of food addiction-like behavior (chocolate-flavoured pellets across FR1 and FR5 sessions) (Fig 5A), addicted mice showed a higher number of reinforcers earned per session compared with non-addicted mice (mean ± SEM; repeated-measures ANOVA, *P = 0.035) (Fig. 2A). Furthermore, addicted mice displayed higher persistence of responding (*P = 0.021) and higher compulsivity (*P = 0.041) than non-addicted mice. In contrast, both addicted and non-addicted mice showed similar motivation. The distribution of addiction-like phenotypes based on three behavioural criteria yielded 3 addicted and 12 non-addicted mice, according to thresholds set at the 75th percentile of the whole cohort (Fig. 2E).

**Figure 2.**
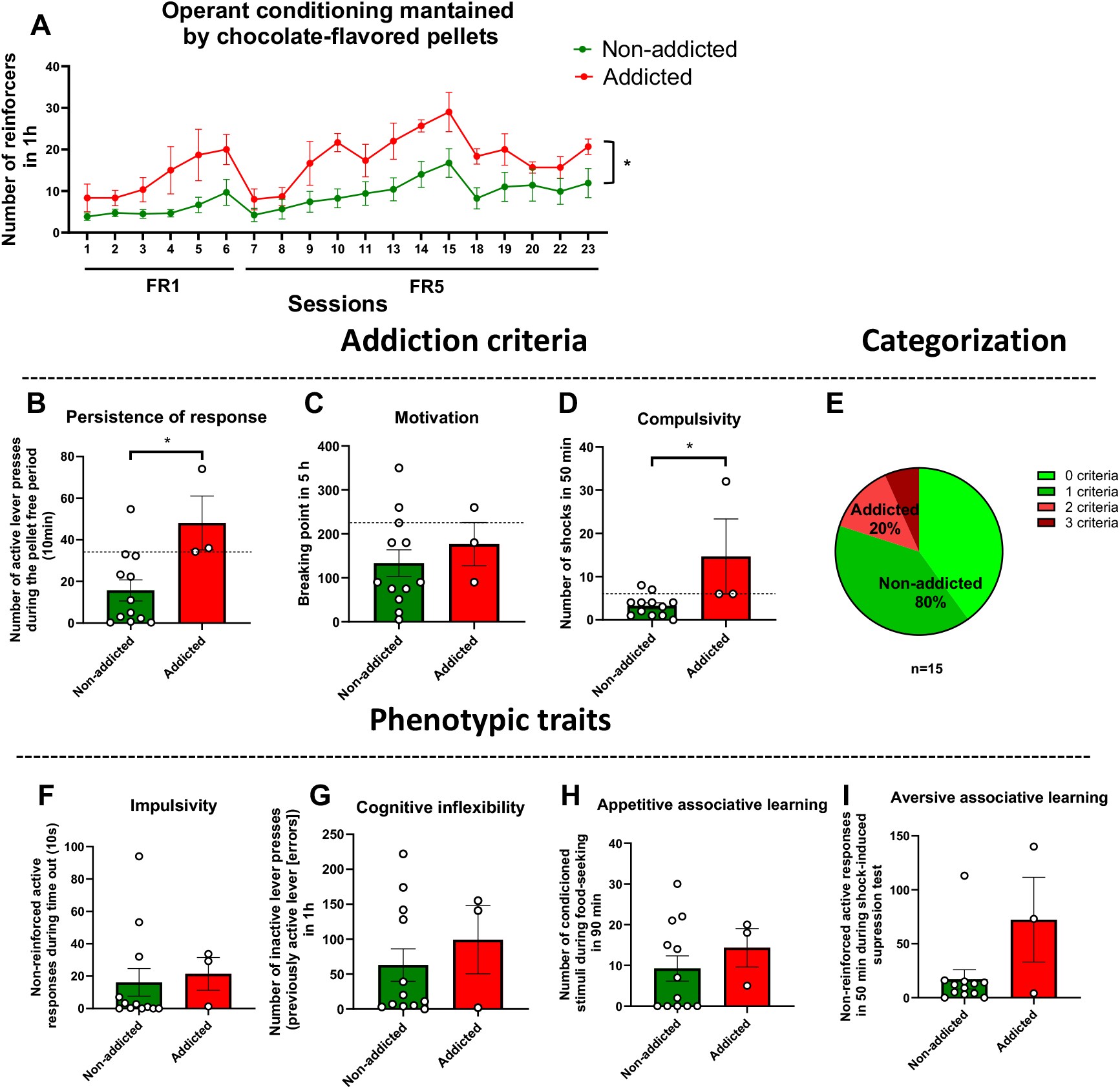
Assessment of food addiction-like behaviour. (A) Operant responding maintained by chocolate-flavoured pellets across FR1 and FR5 sessions (mean ± SEM; repeated-measures ANOVA, *P = 0.035). (B–D) Addiction-like criteria: (B) Persistence of response (Mann–Whitney U test, *P = 0.021). (C) Motivation. (D) Compulsivity (Mann–Whitney U test, *P = 0.041). The dashed horizontal lines represent the 75th percentile of the entire cohort and were used as thresholds to classify mice as positive for each criterion. (E) Distribution of addiction-like phenotypes based on the three behavioural criteria (n = 3 addicted; n = 12 non-addicted). (F–I) Behavioural traits associated with vulnerability to addiction: (F) Impulsivity. (G) Cognitive inflexibility. (H) Appetitive associative learning. (I) Aversive associative learning.

Additional behavioral traits associated with vulnerability to addiction were also evaluated (Fig. 2B–D). No significant differences were observed between addicted and non-addicted mice in motor impulsivity, cognitive inflexibility, aversive associative learning, and appetitive associative learning (Fig. 2F-I).

During specific behavioral events, PL mPFC activity was consistently lower in addicted mice than in non-addicted mice (Fig. 3). In the persistence of response test, addicted mice exhibited significantly reduced normalized PL global fluorescence during active lever presses in the pellet-free period (***P < 0.001) (Fig. 3A). No significant differences were observed in motivation (Fig. 3B). In the compulsivity test, addicted mice showed significantly reduced PL activity when all shocks were considered (*P = 0.013, Fig. 3C). This decreased activity was also evident when the analysis was restricted to the 5th shock (**P = 0.007). Addicted mice further showed reduced PL mPFC activity than non-addicted mice during premature responses in the time-out period (impulsivity; P = 0.039, Fig. 3D), during inactive lever presses in the reversal learning session (cognitive inflexibility; **P < 0.001, Fig. 3E), and during active lever presses in the post-shock session (aversive associative learning; ***P < 0.001, Fig. 3F).

**Figure 3.**
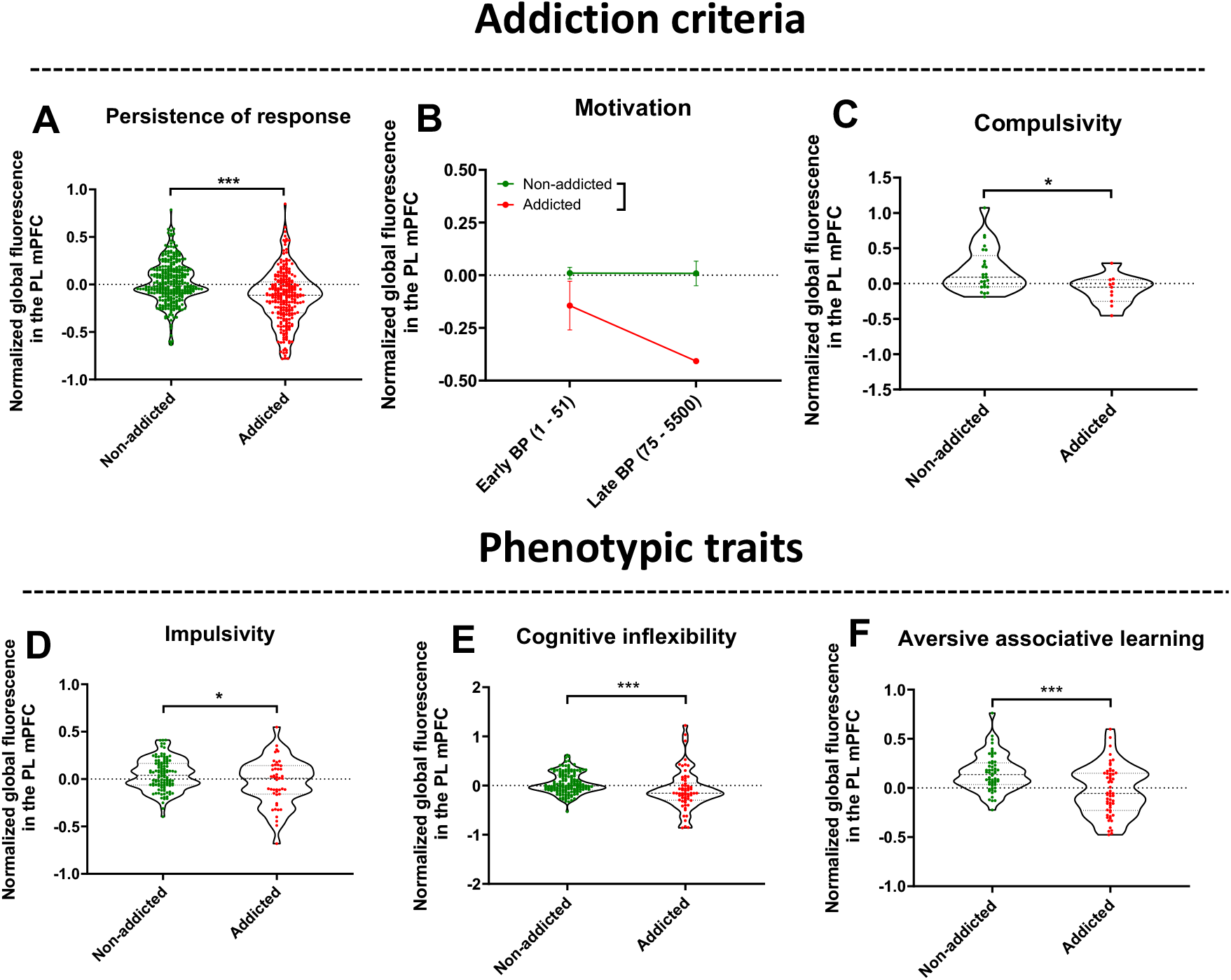
Evaluation of prelimbic (PL) medial prefrontal cortex (mPFC) activity during specific behavioural events. (A) Persistence of response: Normalized PL mPFC mglobal fluorescence during active lever presses in the pellet-free period (PFP) (Mann–Whitney U test, ***P < 0.001). (B) Motivation: Normalized PL mPFC global fluorescence when surpassing early (1–51) and late (75–5500) breaking points during the progressive ratio session. (C) Compulsivity (all shocks): Normalized PL mPFC global fluorescence aligned to shock delivery during the compulsivity test (Mann–Whitney U test, *P = 0.013). (D) Impulsivity: Normalized PL mPFC global fluorescence during premature active responses in the time-out period (TO) (Student’s t test, P = 0.039). (E) Cognitive inflexibility: Normalized PL mPFC global fluorescence during inactive lever presses (errors) in the reversal learning session (Mann–Whitney U test, ***P < 0.001). (F) Aversive associative learning: Normalized PL mPFC global fluorescence during active lever presses in the post-shock session (Student’s t test, ***P < 0.001).

At the single-neuron level, the proportion of event-responsive PL mPFC neurons differed between addicted and non-addicted mice across several behavioral domains (Fig. 4). The percentages of neurons classified as excited or inhibited were quantified during reinforcer delivery across FR1–FR5 sessions (Fig. 4A), persistence of respons (Fig. 4B), motivationally relevant events in the progressive ratio task (Fig. 4C), shock delivery (all shocks and 5th shock only, Fig. 4D), premature responses in the time-out period (Fig. 4E), cognitive inflexibility (Fig. 4F), and active lever presses in the post-shock session (Fig. 4G). Addicted mice showed a significantly higher proportion of inhibited PL mPFC neurons during active lever presses in the pellet-free period compared with non-addicted mice, a behavioral response that mimic motor impulsivity (***P < 0.001, Fig. 4B). During inactive lever presses (errors) in the reversal session of cognitive inflexibility, addicted mice exhibited a significantly lower proportion of excited PL neurons than non-addicted mice (*P = 0.003, Fig. 3F).

**Figure 4.**
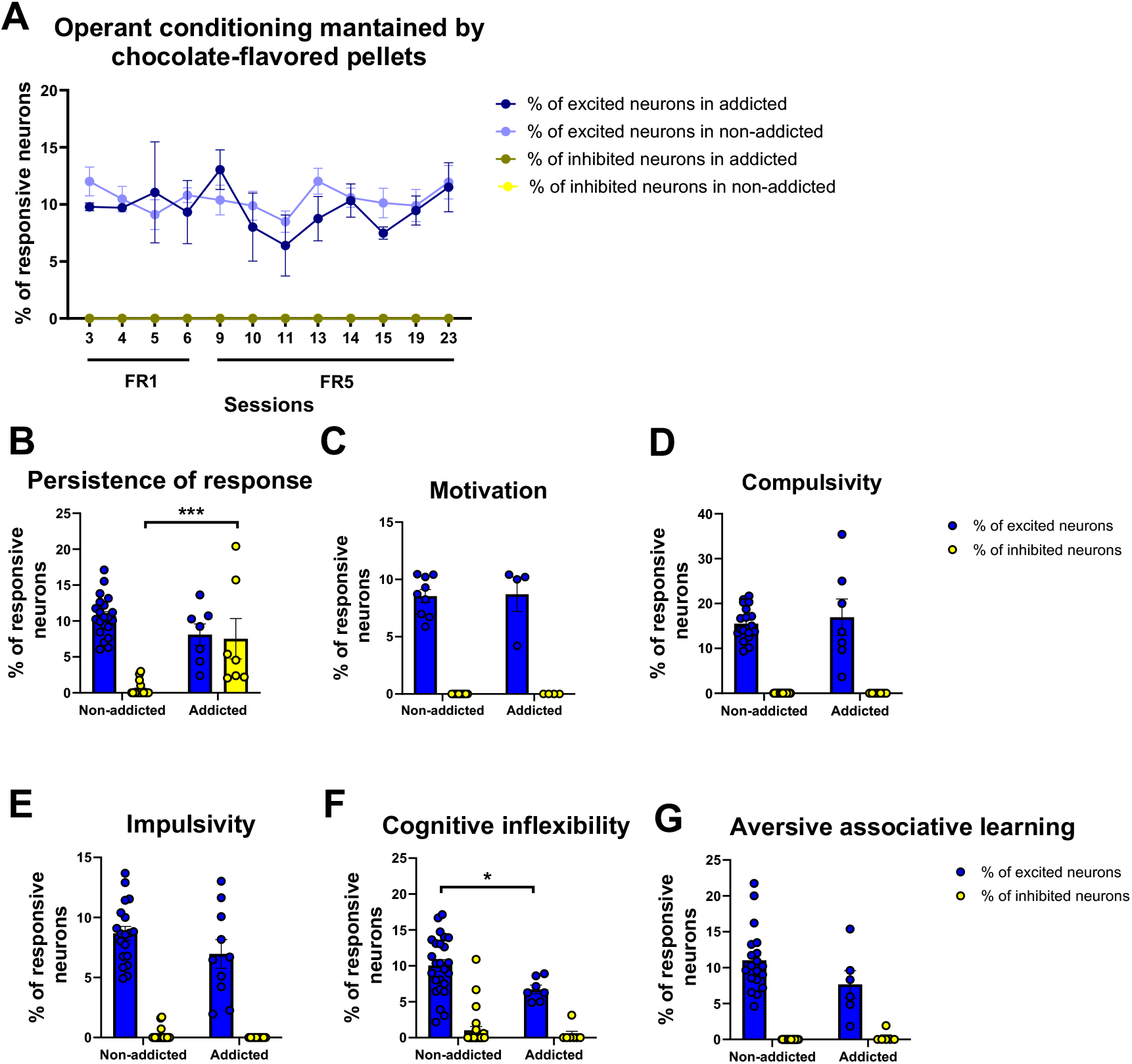
Identification of event-responsive neurons. (A) Percentage of PL mPFC neurons classified as excited or inhibited during reinforcer delivery across operant conditioning sessions (FR1–FR5). (B) Persistence of response: Percentage of PL mPFC neurons excited or inhibited during active lever presses in the pellet-free period (PFP) (Mann–Whitney U test, ***P < 0.001). (C) Motivation: Percentage of responsive neurons during events in which mice surpassed early, late, or any breaking point in the progressive ratio session. (D) Compulsivity (all shocks): Percentage of neurons responsive to shock delivery during the compulsivity test. (E) Impulsivity: Percentage of PL mPFC neurons responsive during premature active responses in the time-out period (TO). (F) Cognitive inflexibility: Percentage of PL mPFC neurons responsive during inactive lever presses (errors) in the reversal session (Student’s t test, *P = 0.003). (G) Aversive associative learning: Percentage of PL mPFC neurons responsive during active lever presses in the post-shock session.

Verification experiments confirmed AAV expression and GRIN lens placement within the PL mPFC, with viral spread and lens tracts localized to the PL region at stereotaxic coordinates AP + 1.8 mm, LM ± 0.3 mm, DV –2.3 mm (Fig. S1). This histological verification supports that the recorded signals reflect PL neuronal activity.

## DISCUSION

We identified differential PL mPFC calcium-dynamics signature in mice depending on the behavioral phenotype, revealing marked differences between addicted and non-addicted mice following operant training in a food addiction model [16]. Notably, mice that met food addiction criteria exhibited reduced calcium activity during persistence of response and compulsivity— two behavioral domains strongly associated with PL mPFC function. In contrast, no differences between addicted and non-addicted animals emerged in motivation, a phenotype more closely linked to nucleus accumbens circuitry. Our findings demonstrate that PL mPFC activity is consistently attenuated in food-addicted mice across multiple behavioral domains, supporting a model in which diminished prefrontal engagement contributes to the emergence and expression of addictive-like behaviors.

Addicted mice showed markedly reduced PL calcium activity during persistence of responding, particularly in the pellet-free period, a phase requiring behavioral inhibition and top-down control. This reduction aligns with previous evidence implicating PL dysfunction in impaired inhibitory control [7,8], a hallmark of compulsive reward seeking. In contrast, PL activity during motivation was unaffected, consistent with the idea that motivational drive in food addiction relies more on subcortical structures, such as the nucleus accumbens, rather than on prefrontal cortex areas.

The reduced PL responses observed during compulsivity further highlight the role of mPFC hypofunction in the persistence of maladaptive behaviors despite adverse consequences. Addicted mice exhibited suppressed PL activity both across all shock events and specifically during the 5th shock, suggesting that deficits in processing aversive feedback or in updating action–outcome contingencies may underlie this compulsive-like responding. Similarly, the lower PL activity observed during motor impulsivity tasks and reversal-learning errors indicates broader impairments in impulse control and cognitive flexibility—two executive functions strongly governed by the mPFC [17]. The diminished PL engagement during aversive associative learning also suggests that addicted mice may be less able to integrate aversive experiences into adaptive behavioral strategies.

Single-neuron analyses revealed complementary changes in the distribution of responsive PL mPFC neurons. Addicted mice displayed a significantly higher proportion of inhibited neurons during persistence of responding, indicating a shift in the overall excitatory–inhibitory balance of PL circuit dynamics during a pellet-free period. Across tasks assessing reward processing, persistence, punishment sensitivity, impulsivity, and flexibility, the altered proportions of excited and inhibited neurons suggest that addictive behavior is associated with widespread remapping of PL responsivity. Notably, during cognitive inflexibility, addicted mice showed decreased proportion of excited neurons during erroneous inactive lever presses, potentially reflecting maladaptive or noisy encoding of incorrect action strategies.

Together, these neural activity patterns point to a consistent signature in the PL mPFC specific for mice developing food addiction. Indeed, food addiction was accompanied by a blunted or dysregulated PL mPFC response during behaviors that require executive control, behavioral adaptation, and the integration of aversive feedback. Such prefrontal cortex dysfunction may compromise the ability to suppress inappropriate reward-seeking behavior, thereby facilitating the development and maintenance of compulsive consumption, and ultimately, the loss of eating control. In conclusión, these results pave the way to elucidating how prefrontal circuit dysfunction contributes to loss of eating control and may guide the development of circuit-based strategies to mitigate addictive-like behaviors.

## Acknowledgments

We thank M. Linares, R. Martín, D. Real, F. Porrón, for their technical support. We thank T. Gusinkaia for her bioinformatic support.

## Author contributions

E.M-G. and R.M. conceived and designed the experimental approaches in animal studies coupled with calcium imaging, with the support of R.A. R.C., and I.M-B. performed the behavioral phenotype characterization and the surgeries. C. I. and J. P. performed the bioinformatic analyses. E.M.-G. and R.M. wrote the manuscript, R.C. and I. M.-B. prepared the figures and wrote the results and methods with the support of C. I. and J. P., and the critical review of all authors.

## Data availability

All data are available in the main text or supplementary materials. Correspondence and requests for materials should be addressed to Elena Martín-García.

## Ethics Statement

Animal procedures were conducted in strict accordance with the guidelines of the European Communities Council Directive 2010/63/E.U. and approved by the local ethical committee (Comitè Ètic d’Experimentació Animal-Parc de Recerca Biomèdica de Barcelona, CEEA-PRBB, agreement N°9213). In agreement, maximal efforts were made to reduce the suffering and the number of mice used.

## Supplementary Materials

Fig. S1

## Figures and legends

**Figure S1.**
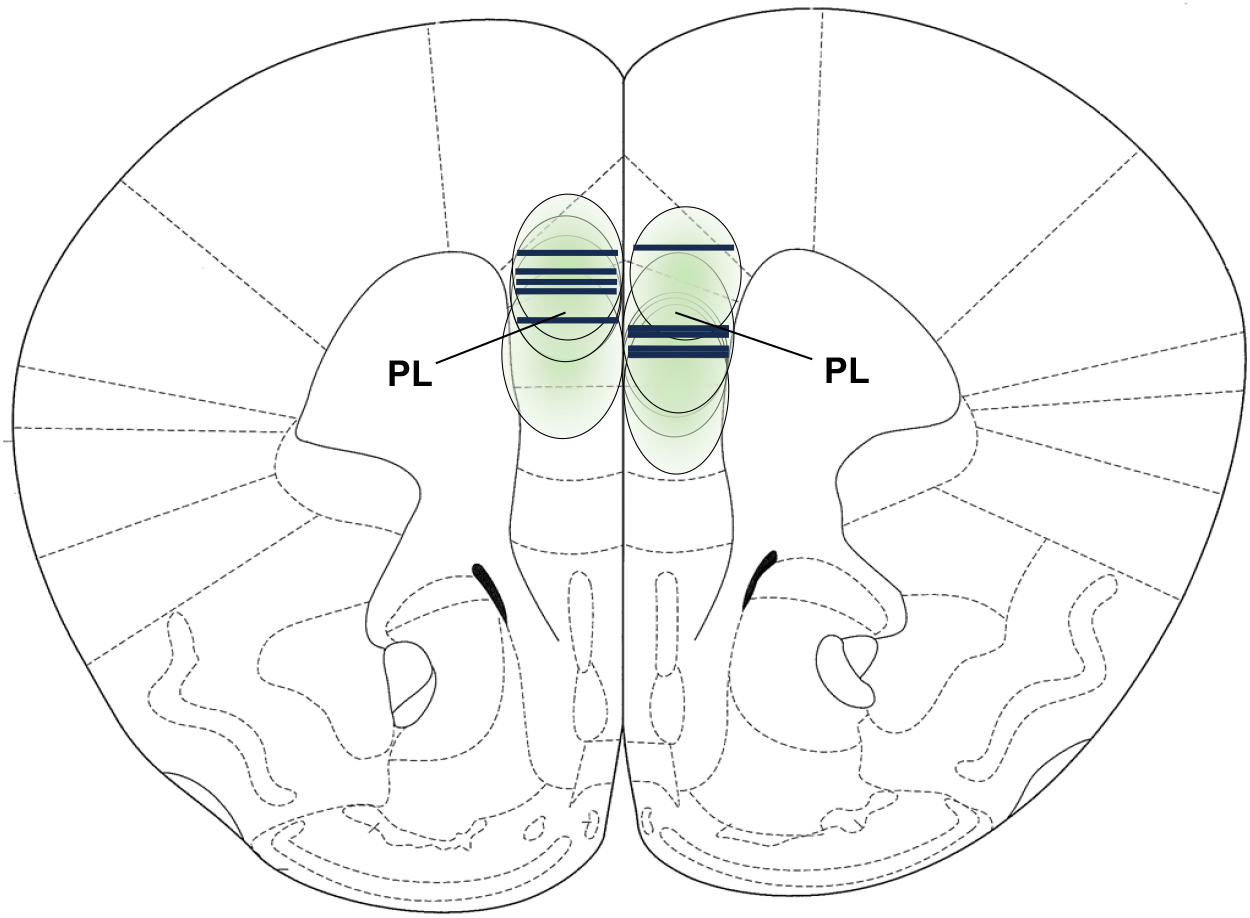
Verification of AAV expression and GRIN lens placement in the prelimbic (PL) medial prefrontal cortex (mPFC). Mouse brain atlas image showing AAV vector spread (outlined green colour) and GRIN lens placement (black bars) in the PL mPFC region. Stereotaxic coordinates for lens placement: AP + 1.8 mm, LM ± 0.3 mm, DV –2.3 mm. Adapted from Paxinos & Franklin’s Mouse Brain Atlas (2008).

